# From birds to mammals: spillover of highly pathogenic avian influenza H5N1 virus to dairy cattle led to efficient intra- and interspecies transmission

**DOI:** 10.1101/2024.05.22.595317

**Authors:** Leonardo C. Caserta, Elisha A. Frye, Salman L. Butt, Melissa Laverack, Mohammed Nooruzzaman, Lina M. Covaleda, Alexis C. Thompson, Melanie Prarat Koscielny, Brittany Cronk, Ashley Johnson, Katie Kleinhenz, Erin E. Edwards, Gabriel Gomez, Gavin Hitchener, Mathias Martins, Darrell R. Kapczynski, David L. Suarez, Ellen Ruth Alexander Morris, Terry Hensley, John S. Beeby, Manigandan Lejeune, Amy K. Swinford, François Elvinger, Kiril M. Dimitrov, Diego G. Diel

## Abstract

Infections with the highly pathogenic avian influenza (HPAI) H5N1 clade 2.3.4.4b virus have resulted in the death of millions of domestic birds and thousands of wild birds in the U.S. since January, 2022^1–4^ Throughout this outbreak, spillovers of the virus to mammals have been frequently documented^5–12^. Here, we report the detection of HPAI H5N1 virus in dairy cattle herds across several states in the U.S. The affected cows displayed clinical signs encompassing decreased feed intake, altered fecal consistency, respiratory distress, and decreased milk production with abnormal milk. Infectious virus and RNA were consistently detected in milk collected from affected cows. Viral staining in tissues revealed a distinct tropism of the virus for the epithelial cells lining the alveoli of the mammary gland in cows. Analysis of whole genome sequences obtained from dairy cows, birds, domestic cats, and a racoon from affected farms indicated multidirectional interspecies transmissions. Epidemiologic and genomic data revealed efficient cow-to-cow transmission after healthy cows from an affected farm were transported to a premise in a different state. These results demonstrate the transmission of HPAI H5N1 clade 2.3.4.4b virus at a non-traditional interface and to a new and highly relevant livestock species, underscoring the ability of the virus to cross species barriers.

## Introduction

The highly pathogenic avian influenza H5Nx goose/Guangdong lineage, an influenza A virus (IAV) from the family *Orthomyxoviridae*, emerged in China in 1996. The viral lineage initially was detected only in poultry, but as early as 2002 there were detections in wild birds. The virus has frequently reassorted with other wild bird and poultry influenza viruses, with the hemagglutinin gene remaining as the only gene that defines this genetic lineage of viruses, and ongoing antigenic changes have required a classification system with numerous clades and subclades^13^. During the past decade, this HPAI H5 lineage of virus evolved into eight clades (2.3.4.4a-2.3.4.4h) of three main neuraminidase subtypes, N1, N8 and N6. In the last three years, the H5N1 clade 2.3.4.4b has been the predominant subtype causing disease outbreaks globally^14,15^. HPAI H5 viruses have infected multiple avian species and have shown a potential to infect humans and other mammalian species^5–8^. The World Health Organization (WHO) has reported a total of 860 human infections with 454 fatal cases since 2003, but this represents a case fatality rate with serologic evidence of more widespread infection with less severe clinical disease. The potential of human-to-human transmission remains low because the virus currently does not transmit efficiently human to human^16^. The viruses of the H5N1 clade 2.3.3.4b have been widely circulating in migratory wild bird populations in Europe, Africa, and Asia since 2016. The first detection in North America (Canada) dates back to December 2021, which represented the first transatlantic spread of the virus^17^. In January 2022, H5N1 was detected in hunter harvested wild birds in North and South Carolina in the U.S.^18^, and was later found in several wild avian species^1^, followed by a spillover into commercial poultry on February 8, 2022^2^ Since 2022, H5N1 infections have resulted in high morbidity and mortality in domestic poultry and the culling of over 90 million birds in the U.S. alone^16^. The continuous and widespread circulation of this high-consequence panzootic pathogen in the U.S. is of major concern and poses a significant threat to animal and public health.

In addition to devastating consequences to domestic and wild avian species, H5N1 clade 2.3.4.4b spillovers have been detected in 30 mammalian species, including 21 wild mammalian species^9,10,19^. In 2022, in Asia, Europe, and U.S., carnivorous wild mammals (e.g. foxes^6^, bears^11^, cats^20^, and harbor seals^5^) infected with H5N1 presented clinical disease with neurologic signs due to underlying encephalitis. The virus has also reached beyond the North and South polar circles, killing a polar bear in the Arctic and seals and gentoo penguins in Antarctica^21^. In 2022, a H5N1 human infection was reported in Colorado, U.S., causing mild clinical signs in a poultry farm worker who recovered from the infection^22^. During 2023, two major outbreaks of H5N1 in harbor seals resulted in high mortality in Maine and Washington. Two domestic indoor-outdoor cats were acutely neurologic and died from HPAI-induced encephalitis in January 2023 in Nebraska^12^. Acute death of a skunk was reported in Washington in 2023^23^ with several more reports in this species in 2023 and 2024^24^. On March 20, 2024, the Minnesota board of animal health reported that a juvenile goat tested positive for HPAI, representing the first report of HPAI H5N1 infection in a livestock species; backyard poultry had previously tested positive on the same premises^25^.

Here we report the spillover of HPAI H5N1 clade 2.3.4.4b virus into dairy cattle and describe the findings of a clinical, pathological, and epidemiological investigation including genomic characterization of the viruses obtained from infected dairy cattle, wild birds, and other mammals from nine affected farms (Farms 1 to 9) across four states in the U.S.

## Results

### Clinical and epidemiological investigation

In February-March 2024, a morbidity event of unknown etiology affecting dairy cattle was reported in several farms in Texas (TX), New Mexico (NM), and Kansas (KS), with subsequent spread to other states. We conducted a clinical and epidemiological investigation in nine affected farms, 8 located in the southwestern U.S., including in TX (Farms 1, 2, 5, 6, and 7), NM (Farms 4 and 8), and KS (Farm 9), and one farm in OH (Farm 3), which was affected after apparently healthy lactating cattle were moved from TX to this location. Affected dairy cattle in these farms presented with decreased feed intake, decreased rumination time, mild respiratory signs (clear nasal discharge, increased respiratory rate, and labored breathing), lethargy, dehydration, dry/tacky feces or diarrhea, and milk with abnormal yellowish colostrum-like color, thick and sometimes curdled consistency. Additionally, an abrupt drop in milk production, with several affected animals presenting no milk secretion, was noted. Upon clinical examination, mammary gland involution was observed in several of the affected cows (**Extended Data Fig. 1**). The rate of clinically affected animals ranged between 3% and 20%. Notably, several of the affected farms reported simultaneous mortality events in wild birds (great-tailed grackles), peridomestic birds (pigeons), and in outdoor domestic (cats) and wild mammals (raccoons) (**Extended Data Table 1**). The clinical disease in dairy cattle lasted 5-14 days, with animals returning to pre-outbreak health status, rumination times, and feed intake, but maintaining decreased milk production for at least four weeks.

### Detection of HPAI H5N1 in dairy cattle, birds, and other mammals

A broad diagnostic investigation was conducted in samples collected from Farms 1-9. Initially, nasal swabs, serum, and blood buffy coats from 10 affected cows from Farm 1 were subjected to viral metagenomic sequencing. Influenza A virus sequences were obtained from one nasal swab. Real-time reverse-transcriptase PCR (rRT-PCR) targeting the IAV matrix (M) and hemagglutinin 5 (H5) genes confirmed H5 IAV infection. Eight out of ten paired milk samples collected from the same cows also tested positive for HPAI-H5N1 virus (**Supplementary Data Table 1**). Additionally, oropharyngeal swabs from great-tailed grackles and pigeons, and lung and brain tissues from a cat found dead on Farm 1 tested positive for HPAI-H5N1 clade 2.3.4.4b virus RNA by rRT-PCR (**Supplementary Data Table 1**).

A similar epidemiological scenario involving mortality events in domestic and wild mammals was observed in Farms 3, 4, 5, and 8. Six domestic cats died in Farm 3 after the disease onset in dairy cows. Cats found dead on Farms 4 and 5 and cats and a raccoon found dead on Farm 8 tested positive for HPAI H5N1 via rRT-PCR. In all occurrences, these animals died after the onset of the clinical outbreak in dairy cattle.

Testing of multiple sample types (n=331) collected from cattle from Farms 1-9 by rRT-PCR showed viral RNA detection sporadically in nasal swabs (10/47), whole blood (3/25), and serum (1/15), and most frequently in milk samples (129/192). The milk samples consistently had the highest viral RNA loads by rRT-PCR of the samples tested. (**Fig. 1A; Supplementary Data Table 1**). Results from rRT-PCR performed on tissues collected from three affected cows revealed the presence of viral RNA in lymph nodes, lung, small intestine, and mammary gland. The highest viral RNA loads were detected in the mammary gland (**Fig. 1B; Supplementary Data Table 1**), supporting the results showing high viral load and shedding in milk. Additionally, hemagglutination inhibition antibody testing in paired serum samples collected from animals (n=20) in Farm 2 confirmed H5N1 infection in affected dairy cows (**Fig. 1C**).

**Figure 1.**
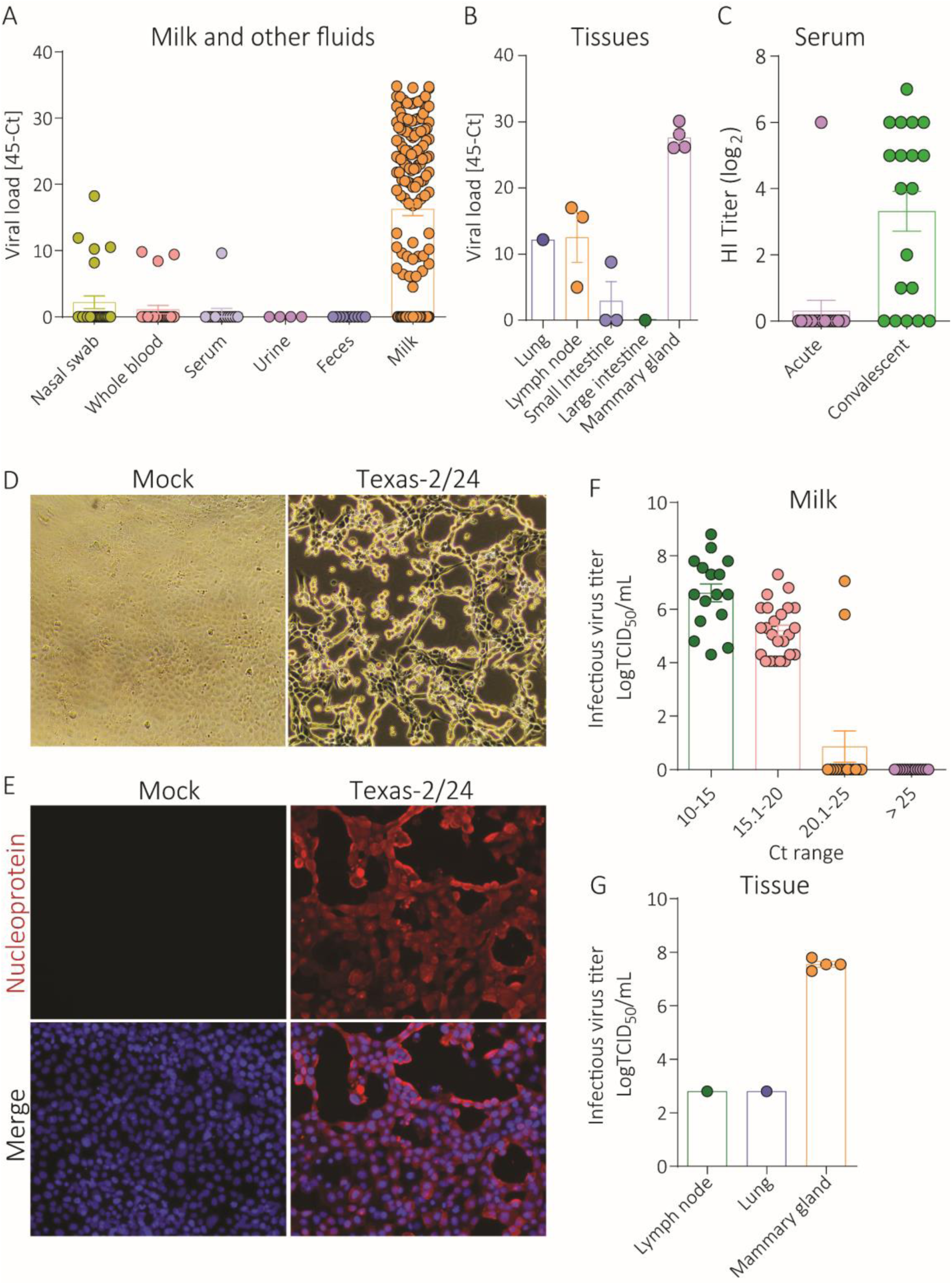
Detection and isolation of HPAI H5N1 in dairy cattle. A) Viral RNA loads in nasal swab, whole blood, serum, urine, feces, and milk samples collected from cattle from Farms 1-9 quantified by rRT-PCR targeting the influenza A virus matrix gene. B) Viral RNA loads in tissues of dairy cattle quantified by rRT-PCR targeting the influenza A virus matrix gene. C) Serum antibody responses in affected cattle quantified by hemagglutination inhibition (HI) assay. (D) Cytopathic effect of HPAI H5N1 virus from milk in bovine uterine epithelial cells Cal-1. E) Detection of infectious HPAI virus in Cal-1 cells by immunofluorescence assay using a nucleoprotein specific monoclonal antibody (red) counterstained stained with 4′,6-diamidino-2-phenylindole (Blue). Infectious HPAI virus in milk (F) and tissues (G) detected by virus titration. Virus titers were determined using endpoint dilutions and expressed as TCID_50_.mL^-1^. The limit of detection (LOD) for infectious virus titration was 10^1.05^ TCID_50_.mL^-1^.

### Infectious HPAI virus shedding in dairy cows

Virus isolation and viral quantifications were performed on milk samples from Farms 1, 2, and 3. Infectious HPAI H5N1 virus was isolated in bovine uterine epithelial cells (Cal-1) from pooled milk pellet samples from 10 cows from Farms 1 and 2 (**Fig. 1D-E**). Notably, virus titers in milk from affected animals ranged from 10^4.0^ to 10^8.8^ 50% tissue culture infectious dose (TCID_50_) per ml (**Fig. 1F**), demonstrating efficient shedding and high viral load in milk from infected animals. A high viral load ranging from 10^7.3^ to 10^7.8^ TCID_50_.ml^-1^ was also detected in mammary gland tissues (**Fig. 1G**).

### Shedding in clinical versus non-clinical cattle

Virus shedding was investigated in samples (milk, nasal swabs, urine, and feces) collected from clinical and non-clinical animals from Farm 3. Overall, virus shedding was detected more frequently in milk samples from clinical animals (24/25) with higher RNA viral load compared to non-clinical animals (1/15) (**Fig. 2A**, **Extended Data Table 2**). Clinical animals shed virus at a lower frequency in nasal swabs (6/25) and urine (2/15), and no viral RNA was detected in feces. (**Fig. 2A**, **Extended Data Table 2**). In non-clinical animals, viral RNA was detected in 6/19 nasal swabs and 4/8 urine samples (**Fig. 2A**, **Extended Data Table 2**) indicating subclinical infection.

**Figure 2.**
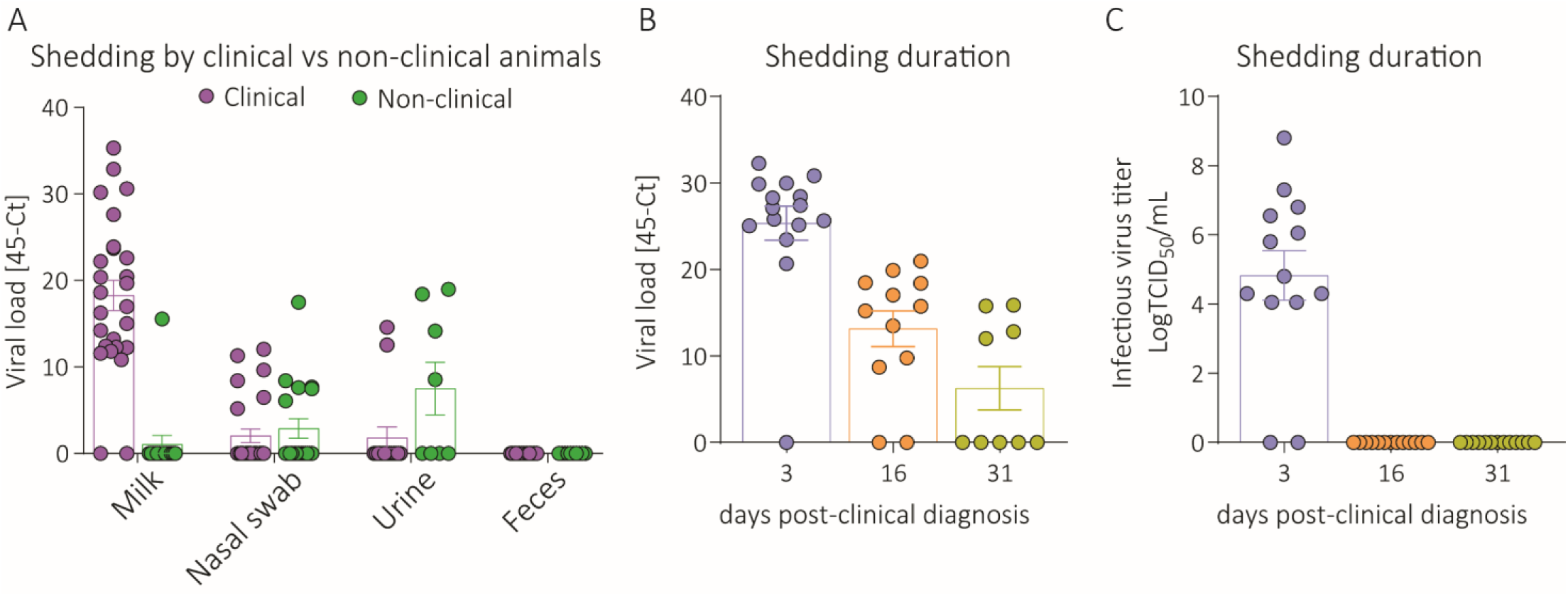
Virus shedding patterns. A) Viral shedding and RNA load in milk, nasal swabs, urine and feces collected from clinical and non-clinical animals from an HPAI affected farm. B) Viral RNA loads in milk samples collected from cattle from Farm 3 on days 3, 16 and 31 post-clinical diagnosis quantified by rRT-PCR targeting the influenza A virus matrix gene. C) Infectious HPAI virus in milk detected by virus titration. Virus titers were determined using endpoint dilutions and expressed as TCID_50_.mL^-1^. The limit of detection (LOD) for infectious virus titration was 10^1.05^ TCID_50_.mL^-1^. The limit of detection (LOD) for infectious virus titration was 10^1.05^ TCID_50_.mL^-1^.

### Duration of HPAI virus shedding in affected dairy cattle

Paired nasal swabs, whole blood, serum, and milk samples were collected at ∼3 (n=15), 16 (n=12), and 31 (n=12) days post-clinical diagnosis pcd of HPAI to assess duration of virus shedding. On day 3 viral RNA was detected in nasal swabs from 2/15 animals, in whole blood of 1/15 animals, in serum of 1/15 animals, and in milk of 14/15 animals (**Fig. 2B, Supplementary Data Table 2**). Importantly, while no HPAI virus RNA was detected in nasal swabs, whole blood, or serum samples collected on days 16 and 31 pcd, 10/12 animals tested on day 16 and 4/12 animals tested on day 31 pcd still had detectable viral RNA in milk (**Fig. 2B, Supplementary Data Table 2**). High infectious viral loads were detected in milk samples on day 3 pcd (10^4.05^ to 10^8.80^ TCID_50_/mL), while no infectious virus was recovered from samples from day 16 and 31 pcd (**Fig. 2C**).

### HPAI virus presents preferential tropism to the mammary gland tissue of infected cows

Histological examination of tissues from affected dairy cows revealed marked changes consisting of neutrophilic and lymphoplasmacytic mastitis with prominent effacement of tubuloacinar gland architecture filled with neutrophils admixed with cellular debris in multiple lobules in the mammary gland (**Fig. 3A**). The most pronounced histological changes in the cat tissues consisted of mild to moderate multi-focal lymphohistiocytic meningoencephalitis with multifocal areas of parenchymal and neuronal necrosis (**Fig. 3B**). A summary of the histologic features observed on a full set of tissues is presented in **Extended Data Fig. 2**.

**Figure 3.**
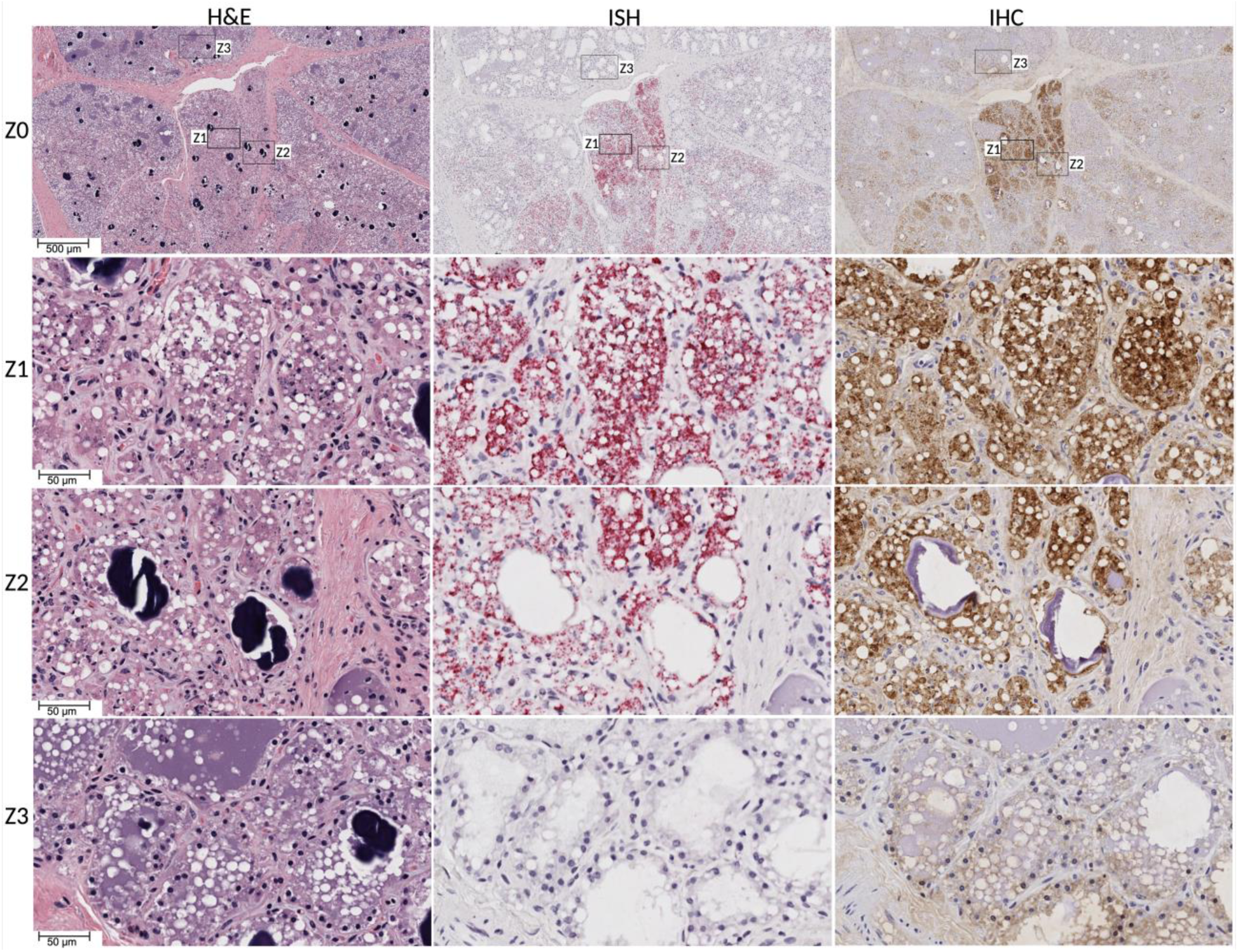
Highly pathogenic avian influenza virus H5N1 detection in dairy cattle mammary gland tissue. Hematoxylin and eosin (H&E) staining (left panels) showing intraluminal epithelial sloughing and cellular debris in mammary alveoli (Z1 and Z2). Normal mammary alveoli filled with milk and fat globules (Z3). In situ hybridization (ISH) (middle panels) targeting Influenza A virus (Matrix gene) showing extensive viral RNA in milk-secreting epithelial cells in the alveoli and in intraluminal cellular debris (Z1 and Z2). Normal mammary alveoli showing no viral staining (Z3). Immunohistochemistry (IHC) (right panels) targeting Influenza A virus M gene showing intracytoplasmic immunolabeling of viral antigen in milk secreting alveolar epithelial cells (Z1 and Z2). Normal mammary alveoli showing no viral staining (Z3)

Using in situ hybridization (ISH) and immunohistochemistry (IHC), viral RNA and antigen were detected in mammary gland, lymph node, trachea (negative with IHC), spleen, colon and heart of affected cattle and in the brain (cerebrum, cerebellum, and brain stem), lung and liver of the affected cat (**Extended Data Table 3**, **Extended Data Fig. 3**). Viral RNA and antigen were detected in the nucleus and cytoplasm of alveolar epithelial cells in mammary gland and peripheral areas of germinal centers of lymph nodes. In mammary glands, viral RNA and antigen was present in the alveolar milk-secreting epithelial cells and inter acinar spaces. In the affected cat, viral RNA and antigen were present in neuronal soma and glial cells in the brain and reticular epithelial cells in the spleen. Viral RNA and protein staining were observed in cerebral neurons, glial cells, endothelial cells lining the choroid plexus, and Purkinje cells in the molecular layer of cerebellum. In lung, viral RNA labeling and immunoreactivity was observed in bronchiolar epithelial cells and alveolar type II pneumocytes (**Extended Data Fig. 2A-2B, Extended Data Table 3**). These results demonstrate a distinct tropism of HPAI H5N1 virus for the mammary tissue of cattle and the central nervous system tissue of cats.

### Reassortment event preceded spillover of HPAI H5N1 virus into dairy cattle

All sequences obtained from the farms in our study (n=91) were classified within a newly emerging B3.13 genotype, which comprises PA, HA, NA and M gene segments of an Eurasian wild bird ancestry (ea1), while PB2, PB1, NP and NS gene segments originated from American bird lineages (am1.1, am2.2, am4, and am8, respectively) (**Extended Data Table 4**). To identify parent genotypes and to define the approximate timeline of the reassortment events that led to the emergence of genotype B3.13, we analyzed the influenza sequence database focusing on sequences obtained during 2021-2024. The flow of gene segments suggests that the B3.13 genotype is a reassortant virus that acquired PB2 and NP gene fragments from low pathogenic avian influenza (LPAI) viruses of subtypes H3 and H11 (**Extended Data Table 4**). The first genome segment derived from LPAI American bird lineage to be incorporated in HPAI H5N1 clade 2.3.4.4b was the NS gene (am1.1); the earliest evidence of its emergence derived from a reassortant virus classified as genotype B3.2 obtained from a skunk in Idaho in November of 2022 (A/skunk/Idaho/22-023547-001-original/2022 (H5N1); **Extended Data Table 4**). Incorporation of the am4 PB1 gene segment into HPAI H5N1 clade 2.3.4.4b was first detected in December 2023, in a sequence classified as genotype Minor60 obtained from a Ross goose collected in Kansas (A/Ross’s goose/Kansas/W23-949A/2023 (H5N1). The HPAI H5N1 clade 2.3.4.4b reassortant genotype B3.13 virus, which incorporated the am2.2 PB2 and am8 NP genome segments, was first detected on January 25, 2024 in a Canadian goose in Wyoming (A/Canada goose/Wyoming/24-003692-001-original/2024 (H5N1), followed by a detection in a peregrine falcon in California (A/peregrine falcon/California/24-005915-001-original/2024 (H5N1)) on February 14, 2024, and shortly after on February 23, 2024 in a skunk in New Mexico (A/skunk/New Mexico/24-006483-001-original/2024 (H5N1)). Of note, New Mexico is one of the first states to report clinical outbreaks compatible with HPAI H5N1 infection in cattle. The host species, in which the reassortment event that culminated with the incorporation of the am8 NP segment and the emergence of HPAI H5N1 genotype B3.13 virus, remains unknown.

### Phylogenomic and evolutionary analyses of HPAI H5N1 B3.13 genotype

Phylogenetic analysis revealed that all sequences obtained and related to the dairy cattle outbreaks in Farms 1-9, including sequences obtained from wild birds and mammals on the affected farms, formed a large phylogenetic group (**Extended Data Fig. 2**, **Fig. 4A**), descending from the most recent common ancestral sequence obtained from a skunk in NM on February 23, 2024 (A/skunk/New Mexico/24-006483-001-original/2024). The PB2-based phylogeny revealed that all HPAI H5N1 sequences obtained from the affected dairy farms characterized in the present study formed nine distinct phylogenetic branches. Notably, these phylogenetic groups of closely related sequences were not always formed by sequences derived from the same farm (**Fig. 4B**). A phylogenetic branch formed by sequences from a cat from Farm 5 and cattle sequences from Farms 7 (n=5) and 9 (n=3) suggested a direct epidemiological relationship between these farms (**Fig. 4B**). Similarly, sequences obtained from cattle from Sites 1 and 2 of Farm 2 (a multi-site dairy operation), formed a monophyletic cluster, indicating co-circulation of the virus in these two sites (**Fig. 4B**).

**Figure 4.**
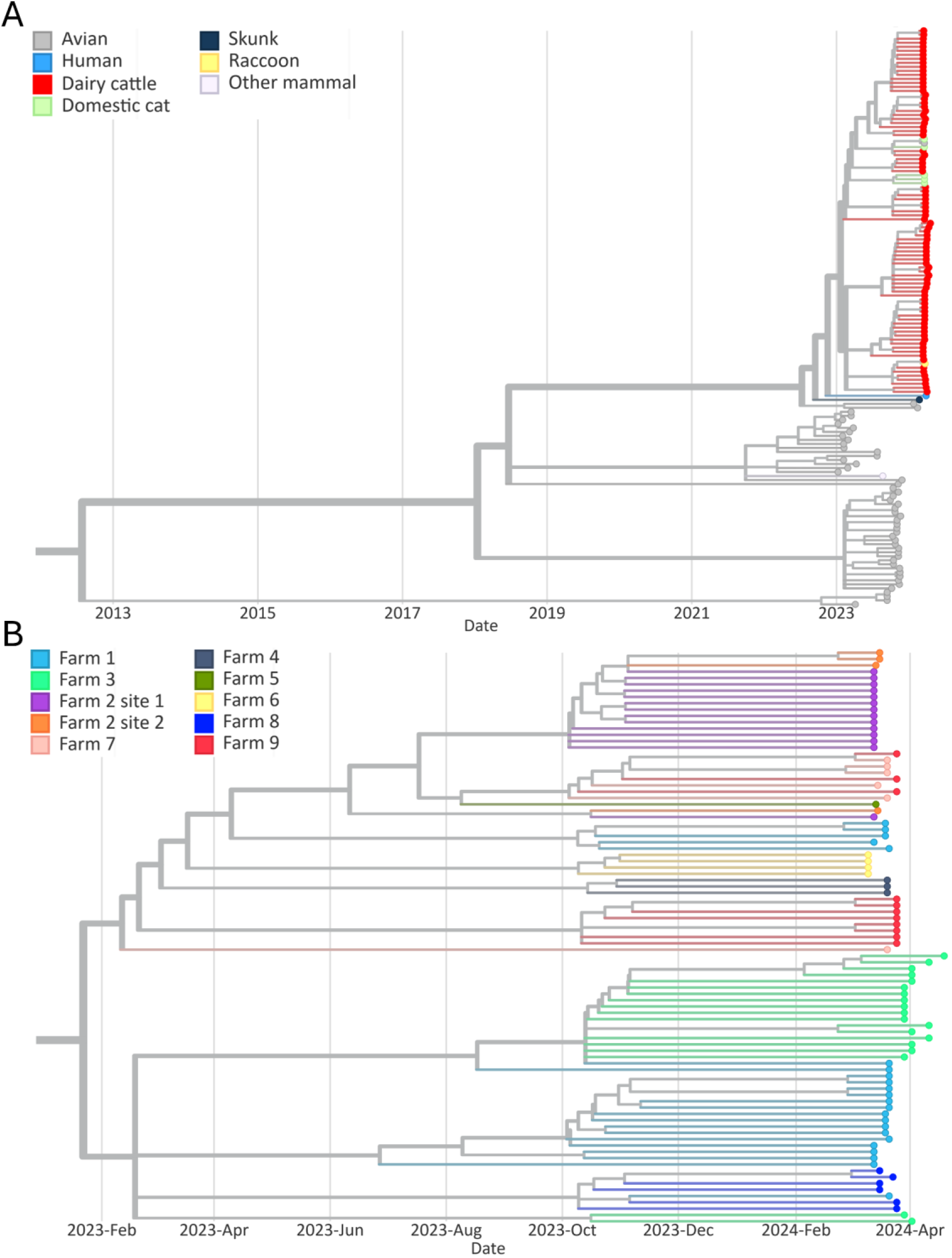
Phylogenetic analysis of the PB2 genome segment. A) Phylogeny of sequences derived from cattle, cats, raccoon, and grackle sampled in the farms described in this study, and other sequences in closer ancestral branches, obtained from GISAID database. Nodes are colored by host species. (B) Detailed view of the clade containing 91 sequences derived from animals sampled in the farms described in this study. Nodes are colored by farm.

Next, the mutation profile of HPAI H5N1 clade 2.3.4.4b was investigated. Initially, we evaluated the occurrence of mutations with known functional relevance to IAV (e.g. host adaptation, virulence, host specificity shift, etc.) in comparison to the original H5N1 A/GsGd/1/1996 virus (**Supplementary Data Table 3**). Further we performed a detailed comparative genome analysis and mutational profiling using sequences obtained in the U.S. throughout the 2021-2024 outbreak (**Extended Data Table 5**). The sequence A/chicken/NL/FAV-0033/2021 2.3.4.4b was used as a reference to identify mutations in different genome segments across species affected by H5N1 clade 2.3.4.4b in the U.S. Representative sequences from multiple genotypes (A1, A2, B1.3, B3.2, Minor01, B3.6, and B3.13) were selected, including sequences from avian (chicken and great tailed grackle) and mammalian (skunk, red fox, harbor seals, human, goat, cat, and cattle) hosts. A total of 132 amino acid substitutions were observed across the 8 genome segments of HPAI H5N1 2.3.4.4b genotype B3.13 (**Extended Data Table 5**). Most of which are low frequency mutations observed in a small proportion of cattle derived viral sequences. Fifteen mutations emerged in viruses circulating in late 2023 and were detected in genotype B3.13 viruses in 2024 including in the PB2 (V109I, V139I, V495I, and V649I), PB1 (E75D, M171V, R430K, and A587P), PA (K113R), HA (T211I), NA (V67I, L269M, V321I, and S339P), NP (S482N), and NS1 (C116S) genes. Seven additional mutations emerged in viruses of genotype B3.13 including five substitutions in PB2 (T58A, E362G, D441N, M631L, and T676A) one in PA (L219I) and one in NS1 (S7L). When compared to the first reported B3.13 sequences (A/falcon/CA/2024(H5N1) and A/skunk/NM/2024(H5N1)), the cattle and domestic cat HPAI H5N1 virus sequences associated with the outbreak in dairy cattle presented five amino acid substitutions, including: three in PB2 (E362G, D441N and M631L), one in PA (L219I) and one in NS (S7L), suggesting that these could have emerged following spillover in these species.

### Dispersal of HPAI H5N1 B3.13 virus between dairy farms

The geographical dispersal dynamics of HPAI H5N1 B3.13 virus between farms was investigated. The HPAI H5N1 genotype B3.13 sequences obtained from farms presenting an epidemiological link (Farm 2: separate production sites [site 1 and 2]; and Farms 1 and 3: animals were transported from Farm 1 to 3) (**Extended Data Table 1**) or presenting closely related viral sequences (Farms 5, 7, and 9) (**Fig. 4B**) were included in our phylogeographic reconstructions (**Fig. 5A**). Haplotype network analysis of the PB2 viral sequences provided support for focusing the dispersal and phylogeographical inferences on Farms 1 and 3, Farm 2, and Farms 5, 7, and 9 (**Fig. 5B**). The phylogenetic relationship and dispersal pathways were inferred based on PB2 gene sequences, the farm location and date of sample collection to reconstruct the dispersal trajectory of the HPAI virus between the farms. The viral sequences recovered from Farm 2, which were collected from two geographically separated production sites (site 1 and 2, approximately 50 Km apart), fell into two phylogenetic clusters, each comprising sequences from both sites, confirming the exchange of the same virus between these premises (**Fig. 5C**). Phylogeographical dispersal analysis of the HPAI H5N1 sequences recovered from Farm 2, point to site 1 as the likely source of the virus, from which it spread to site 2.

**Figure 5.**
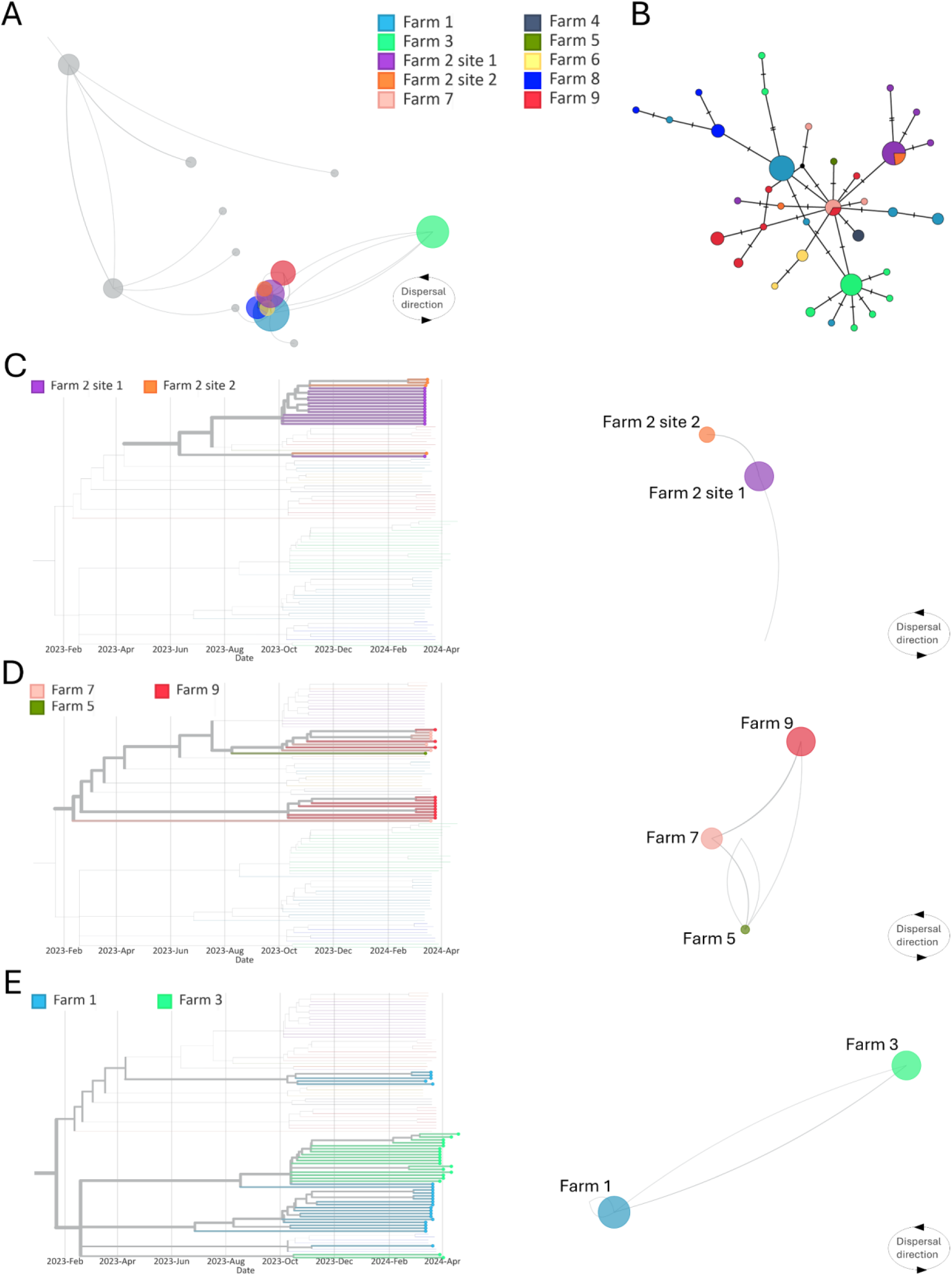
Interstate and local dispersal of HPAI H5N1 genotype B3.13 between farms, based on PB2 gene analysis. (A) Dispersal in North America. Samples described in this study are colored by farm, while locations in grey represent samples from closer ancestral branches obtained from GISAID database. (B) Haplotype network analysis constructed from PB2 segment alignment of HPAI H5N1 obtained from farms described in this study. Different colors indicate different farms. The size of each vertex is relative to the number of samples and the dashes on branches show the number of mutations between nodes. Phylogenetic reconstruction and analysis of dispersal between (C) Sites 1 and 2 of farm 2, (D) Farms 5, 7 and 9, and (E) Farms 1 and 3. Directions of dispersal lines are counterclockwise.

Viral sequences obtained from Farms 5, 7, and 9 (one sequence from cattle in Farm 5, six sequences from Farm 7 and 4 sequences from Farm 9), which are geographically distant from each other (Farm 5 to 7: 186.04 Km, Farm 7 to 9: 280.36 Km; and Farm 5 to 9: 431.01 Km), formed distinct phylogenetic clusters (**Fig. 5D**). A larger cluster comprised one sequence from Farm 5, five sequences from Farm 7, and three sequences from Farm 9, while a smaller and separate branch in the tree was formed by one viral sequence from Farms 7 and eight sequences from Farm 9. Analysis of the directionality of dispersion of HPAIV sequences between these three farms points to Farm 5 as the potential initial source of the virus to Farm 7, from where the virus may have further spread to Farm 9. Given the dispersion of the virus between these three farms, we conducted a broader phylogenetic analysis including other HPAI H5N1 B3.13 sequences available in GISIAD. This analysis revealed two additional H5N1 sequences recovered from blackbirds clustering with Farm 5, 7 and 9 sequences (**Extended Data Fig. 5)**. Worthy of note, the blackbirds were collected between 8-12 Km away from Farm 7. Together these results suggest both long- and close-range lateral spread and transmission of HPAIV between farms.

Sequences obtained from Farms 1 (TX) and 3 (OH) formed five phylogenetic clusters with the largest cluster consisting of two sequences from Farm 1 and 17 sequences from Farm 3. One viral sequence recovered from an animal from Farm 1 was ancestral to all 17 sequences from Farm 3. The second largest cluster was formed mostly by sequences from Farm 1 (n=15). A third smaller cluster comprised five sequences from Farm 1, which were recovered from infected cows, a wild bird, and a domestic cat. Dispersal analysis revealed that HPAIV most likely spread from Farm 1 (TX) to Farm 3 (OH) (**Fig. 4C**). This is consistent with the epidemiological information collected from these farms, which revealed the transportation of 42 apparently healthy dairy cattle from Farm 1 to Farm 3 on March 8, 2024, five days before the first clinical signs were observed in animals in Farm 1 and 12 days before the first clinical animal was identified in Farm 3 (**Extended Data Table 1**). These results indicate transmission of HPAI H5N1 between apparently healthy or subclinically infected dairy cattle.

### Evidence for interspecies transmission of HPAI in affected dairy farms

Given that five of the nine farms included in our study (Farms 1, 3, 4, 5, and 8) reported mortality events in wild (great-tailed grackles) and peri-domestic birds (pigeons), and in wild (raccoon) and domestic mammals (cats), we investigated potential HPAI infection in these species. Whole genome sequencing of the samples from the grackles and a cat from Farm 1 and a raccoon from Farm 8 confirmed infection of these species with a HPAI H5N1 genotype B3.13 virus closely related to the viruses found in dairy cattle in these farms (**Extended Data Fig. 4**). The most recent common ancestor for the cat sequences detected in Farm 1 and the raccoon sequence detected in Farm 8 were obtained from dairy cattle in these farms, indicating cattle-to-cat and cattle-to-raccoon transmission. This is corroborated by epidemiological information revealing that feeding raw milk to farm cats was a common practice in these farms.

## Discussion

Here we describe the spillover of HPAI H5N1 clade 2.3.4.4b virus into dairy cattle and provide evidence of efficient transmission among cattle and between cattle and other species, highlighting the virus’ ability to cross species barriers. This unprecedented spillover and emergence of HPAI H5N1 virus in cattle was preceded by a reassortment event that originated the new genotype (B3.13) comprising genome segments from the Eurasian (ea; PA, HA, NA and M) and American (am; PB2, PB1, NP and NS) bird lineage viruses. The last two genes to be incorporated in the B3.13 virus genome prior to its spillover to cattle were PB2 (am2.2) and NP (am8), suggesting that incorporation of these genes could have resulted in a host range expansion of HPAI H5N1 genotype B3.13. Previous reassortment studies with H5N1 revealed that exchange of the NP gene alone resulted in improved virus replication, expanded tissue tropism, and increased pathogenicity in chickens^26^. In addition to the reassortment event, mutations that appeared following spillover and initial replication of H5N1 genotype B3.13 virus in cattle – for example the PB2 M631L substitution – may have also contributed to enhanced virus fitness in the new cattle host^27^. The PB2 M631L substitution was reported in one study to be critical for increased polymerase activity related to increased severity of disease in mice. Notably, viral sequences obtained from a confirmed human infection with HPAI H5N1 genotype B3.13 in a dairy farm worker in Texas^28^ revealed that the virus did not present the cattle-associated PB2 M631L mutation, but had the well characterized PB2 E627K mutation, related to mammalian adaptation (Ref). Future studies to investigate the function and contribution of the reassorted NP and PB2 genome segments as well as the PB2 M631L mutation on virus host range, tropism, and replication are critical to understand the molecular mechanisms underlying HPAI H5N1 cross species transmission to cattle.

The farms that first reported and confirmed HPAI H5N1 genotype B3.13 infection in cattle in TX, NM, and KS are on the Central North American migratory bird flyway. Importantly, the first reported genotype B3.13 virus genome sequence was obtained from a sample collected from a Canadian goose in Wyoming (January 25, 2024), within the same flyway. This was followed by a detection in a peregrine falcon in California (CA) (February 14, 2024) on the Pacific flyway, with the next reported B3.13 sequence being from a skunk in NM (February 23, 2024), again on the Central flyway. Our modeling using currently available genome sequences suggests dispersal of the newly emerging B3.13 virus from the West (CA) to East (NM and TX), with the sequence recovered from the skunk in NM (A/skunk/New_Mexico/24-006483-001-Original/2024) being the most recent common ancestor to all available sequences from dairy cattle and associated affected species since the onset of the outbreak. The lack of complete epidemiological information regarding the H5N1 genotype B3.13 sequence collected from the skunk in NM precludes definitive conclusions on the link of this animal with affected dairy cattle farms in the region. However, this detection demonstrates the presence of the virus in wildlife in NM around the same time (February-March) in which the first cases of sick cows presenting mild respiratory signs, drop in feed intake, and milk production (which were later confirmed to be caused by HPAI H5N1 genotype B3.13) were reported^29^. Under sampling and limitations of current HPAI wildlife surveillance coupled with lack of diagnostic testing from the initial clinical cases in dairy cattle (likely from early February) complicate investigations on the origin of the outbreak and spillover event into cattle. Obtaining additional historic and prospective sequence data may allow more detailed molecular epidemiological inferences.

Highly pathogenic influenza H5N1 clade 2.3.4.4b infections in mammals have been predominantly associated with neurological invasion and extensive virus replication in the brain^19^. In contrast, the results obtained from our extensive molecular, virological and pathological investigation revealed important and distinct aspects of HPAI H5N1 virus tropism, replication sites, and shedding patterns in dairy cattle. Viral RNA was consistently detected in milk samples, with sporadic detections in nasal swabs and blood (whole blood and serum). Testing of paired nasal swabs, whole blood, serum, urine, and milk samples from affected cattle corroborated these observations and demonstrated consistent virus shedding in milk with sporadic virus detection early in infection (≤5 days pcd) in nasal swabs, whole blood, serum and urine. These results suggested a high tropism of HPAI H5N1 for the mammary gland tissue resulting in a viral-induced mastitis, which was confirmed by histological changes and direct viral staining as well as virus quantification showing high viral loads and replication in the mammary gland. *In situ* hybridization and IHC staining defined the virus tropism and replication to milk-secreting mammary epithelial cells lining the alveoli in the mammary gland. The tropism of HPAI H5N1 for milk-secreting epithelial cells is likely a result of the high expression of sialic acid receptors with an α2,3 (avian-like receptor) and α2,6 (human-like receptor) galactose linkage in these cells^30^. Although the tissue sample size included in our study was small, isolation of the virus in lung and lymph nodes suggests that other organs may also play a role in the virus infection dynamics and pathogenesis in dairy cattle. The initial site of virus replication remains unknown; however, it is possible that the virus can infect through respiratory and/or oral routes replicating at low levels in the upper respiratory tract (e.g. nasal turbinate, trachea, and/or pharynx), from where it disseminates to other organs via a short and low-level viremia. The collected evidence suggests that the mammary gland is the main site of virus replication, resulting in substantial virus shedding in milk. Another possible transmission route includes direct infection of the mammary gland through the teat orifice and cisternae, which could occur mechanically via the milking equipment during milking. In the 1950’s several studies showed that direct inoculation of virus into the udder of dairy cows and goats with the human PR8 strain of type A influenza could result in infection and viral shed for at least 10 days^31–35^. This experimental data suggests, in light of the current outbreak, that mammary epithelial cells, which express α2,3 and α2,6 sialic acid, may be generally susceptible to type A influenza viruses. The uniqueness of this outbreak may be the virus’ likely ability to replicate in other tissues and transmit efficiently. There have been a few other studies that suggest an association between type A influenza and clinical disease^36–41^, but no evidence of sustained transmission or the fulfillment of Koch’s postulates to prove a role in pathogenesis is available. The only published study of goose/Guangdong lineage virus being inoculated into calves show limited viral replication with no clinical disease, but the study was not designed to evaluate transmissibility^42^. Experimental infection studies with HPAI H5N1 genotype B3.13 virus in dairy cattle with sequential and comprehensive sample collections are critical to answer these important questions.

The ability of HPAI H5N1 clade 2.3.4.4b to cross species barriers has been evident and spillover into mammalian species has been reported throughout the current global outbreak^23,43^. Prior to the detection in cattle, however, most mammalian species were considered dead-end hosts, given that virus tropism for the central nervous system commonly resulted in fatal encephalitis^44,45^. Our epidemiological investigation combined with genome sequence- and geographical dispersal analysis provides evidence of efficient intra- and inter-species transmission of HPAI H5N1 genotype B3.13. Soon after apparently healthy lactating cattle were moved from Farm 1 to Farm 3, resident animals in Farm 3 developed clinical signs compatible with HPAI H5N1 providing evidence to suggest that non-clinical animals can spread the virus. Analysis of the genetic relationship between the viruses detected in Farms 1 and 3, combined with phylogeographical modeling indicate that the viruses infecting cattle in these farms are closely related, supporting the direct epidemiological link and indicating long-range viral dispersal and efficient cattle-to-cattle transmission. The results from the phylogenomic and phylogeographical analyses in both sites of Farm 2 and on Farms 5, 6 and 7 also indicate regional long-range farm-to-farm spread of the virus. In these cases, fomites such as shared farm equipment, vehicles, or personnel may have played a role in virus spread. The dispersal of virus between Farms 5, 7 and 9 could have been vectored by wild birds; as suggested by the fact that blackbirds found dead near Farm 7 were infected with a virus closely related to the virus circulating in cattle in these farms. Alternatively, the birds at these premises could have been infected with virus shed by cattle. Our phylogenomic analysis in affected cats (Farms 1, 2, 4, and 5) and the raccoon (Farm 8) combined with epidemiological information revealing the practice of feeding raw milk to cats in these farms indicate cattle-to-cat and cattle-to-raccoon transmission. These observations indicate that complex pathways underlie the introduction and spread of HPAI H5N1 in dairy farms (**Fig. 6**), highlighting the need for efficient biosecurity practices and surveillance efforts in affected and non-affected farms.

**Figure 6.**
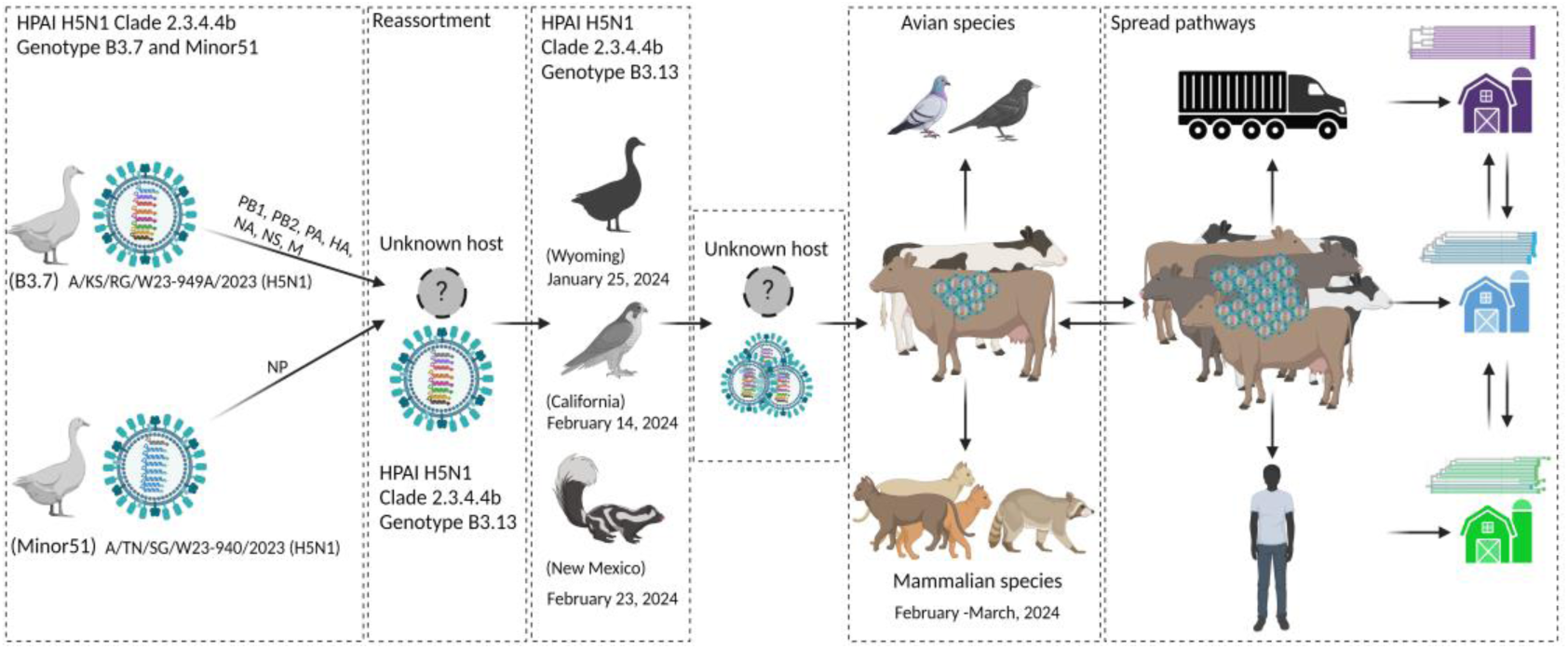
Model of spillover and spread of HPAI H5N1 genotype B3.13 into dairy cattle. A reassortment event in an unknown host species led to the emergence of H5N1 genotype B3.13 which circulated in wild birds and mammals before infecting dairy cattle. Following spillover of H5N1 into dairy cattle, the virus was able to establish infection and efficiently transmit from cow-to-cow (intraspecies transmission) and from cow to other species, including wild (great tailed grackles) and peridomestic birds (pigeons) and mammals (cats and raccoons) (interspecies transmission). Spread of the virus between farms occurred by the movement of cattle between farms, and likely by movement wild birds and fomites including personnel, shared farm equipment and trucks (feed, milk and/or animal trucks).

The spillover of HPAI H5N1 into dairy cattle and evidence for efficient mammal-to-mammal transmission are unprecedented. This newly acquired viral property is concerning as it can lead to adaptation of the virus which may further enhance virus infectivity and transmissibility in other species, including humans. Therefore, it is imperative that robust and continuous surveillance and research efforts be established to monitor the circulation, spread, and adaptation of the HPAI H5N1 virus in this new host species.

## Author contribution

Conceptualization: DGD; Methodology: EF, SLB, ML, MN, LC, ACT, MPK, BC, AJ, KK, ED, GG, GH, MM, ERA, TH; Software: LCC, BK; Validation: SLB, ML, MN, LC, MPK, BC, AJ; Formal analysis: LCC, SLB, ML, BC, DGD; Investigation: LCC, EF, SLB, ML, LC, ACT, MPK, BC, AJ, DRK, MM, ERA; Resources: EF, ACT, DLS, ML, AS, FE, KD, DGD; Data Curation: LCC, EF, SLB, ML, KD, DGD; Writing - Original Draft: LCC, SLB, BK, DGD; Writing - Review & Editing: LCC, EF, SLB, ML, MN, LC, ACT, MPK, BC, AJ, KK, ED, GG, GH, MM, DRK, DLS, ERA, TH, MLV, AS, FE, KD, DGD; Visualization: LCC, SLB, BC, DGD; Supervision: DGD; Project administration: MPK, KD, DGD, Funding acquisition: AS, FE, KD, DGD.

## Data availability

All HPAI H5N1 virus sequences are deposited in GISAID (https:www.gisaid.org/; accession numbers are available in Supplementary Data Table 5), and raw reads have been deposited in NCBI’s Short Read Archive (BioProject number PRJNA1114404). All additional influenza sequences used in our analysis were obtained from GISAID (accession numbers available in Supplementary Data Table 5).

## Acknowledgements

The authors would like to thank the producers and veterinarians who submitted samples and contributed for this investigation. The work was funded by the AHDC, OADDL and TVMDL. This work was supported in part by USDA-NIFA grant no. 2021-68014-33635 (D.G.D, KD) and APHIS NALHN Enhancement grant no. AP21VSD&B000C005 (K.D., D.G.D). We gratefully acknowledge all data contributors, i.e., the Authors and their Originating laboratories responsible for obtaining the specimens, and their Submitting laboratories for generating the genetic sequence and metadata and sharing via the GISAID Initiative, on which this research is based.

## Methods

### Sample collection

Clinical samples used in the present study were collected by field veterinarians from nine clinically affected farms in TX (Farm 1, 2, 4, 5, 6 and 7), NM (Farm 8), KS (Farm 9) or OH (Farm 3). A total of 332 samples collected from dairy cattle (n=323), domestic cats (n=4), great-tailed grackles (n=3), pigeon (n=1) and a racoon (n=1) in the affected farms. All samples including milk (n=211), nasal swabs (n=46), whole blood (n=25), serum (n=15), feces (n=10), urine (n=4), and tissues (mammary gland [n=4], lung [n=1], lymph nodes [n=3], small [n=3] and large intestine [n=1]) from dairy cattle were submitted to the Cornell Animal Health Diagnostic Center (AHDC), Texas A&M Veterinary Medical Diagnostic Laboratory (TVMDL) or the Ohio Animal Disease Diagnostic Laboratory (OADDL) for diagnostic investigations. One domestic cat, two grackles and one pigeon (Farm 1) were submitted to the AHDC while three cats (Farms 4 and 8) and a racoon (Farm 8) and four cows were submitted to TVMDL for necropsy and testing (**Supplementary Data Table 1**).

Sequential samples (milk, nasal swabs and blood) collected from animals (n=15) from Farm 3 were used to investigate duration of virus shedding (**Supplementary Data Table 2**). Additionally, paired samples (milk, nasal sabs, urine and feces) collected from animals presenting respiratory distress, drop in milk production and altered milk characteristics (clinical, n=25) and from apparently healthy animals (non-clinical, n=20) from Farm 3 were used to compare virus shedding by clinical and non-clinical animals (**Extended Data Table 2**).

### Clinical history and epidemiological information

Clinical history from all nine farms were obtained from the sample submission forms sent with the samples to the AHDC, TVMDL and OADDL. Additional relevant information from each farm were obtained from attending veterinarians through investigations conducted by laboratory diagnosticians.

### Real-time reverse transcriptase PCR (rRT-PCR)

Viral nucleic acid was extracted from milk, nasal swabs, whole blood, serum, feces, urine and tissue homogenates. Two hundred µl of milk, nasal swabs, whole blood, serum, and urine were used for extraction. Two hundred µl of raw milk samples were used directly or diluted at the ratio of 1 part of milk to 3 parts of phosphate-buffered saline (PBS) with 200 µl of the dilution used for nucleic acid extraction. Tissues and feces were homogenized in PBS-BSA (1%) (10% w/v), cleared by centrifugation and 200 µl of the supernatant were used for extraction. All RNA extractions were performed using the MagMAX Pathogen RNA/DNA Kit (Thermo Fisher, Waltham, MA, USA) and the automated King Fisher Flex nucleic acid extractor (Thermo Fisher, Waltham, MA, USA) following the manufacturer’s recommendations. The presence of IAV RNA was assessed using the VetMax-Gold AIV Detection Kit (Thermo Fisher, Waltham, MA, USA) and the National Animal Laboratory Network (NAHLN) primers and probe targeting the conserved M gene or the H5 hemagglutinin gene^46^. Amplification and detection were performed using the Applied Biosystems 7500 Fast PCR Detection System (Thermo Fisher, Waltham, MA, USA), under following conditions: 10 min at 45°C for reverse transcription, 10 min at 95 °C for polymerase activation and 45 cycles of 15 s at 94 °C for denaturation and 30 s at 60 °C for annealing and extension. Relative viral loads were calculated and are expressed as 45 rRT-PCR cycles minus the actual CT value (45-Ct). Positive and negative amplification controls as well as internal inhibition controls were run side by side with test samples. Part of samples was also tested using 200 µl of undiluted milk and serum, and 100 µl of whole blood, targeting the M gene. These samples were extracted using the IndiMag Pathogen kit (INDICAL Bioscience) on the KingFisher Flex (Thermo Fisher, Waltham, MA, USA), and the rRT-PCR was performed using the Path-ID™ Multiplex One-Step RT-PCR Kit (Thermo Fisher, Waltham, MA, USA) under following conditions: 10 min at 48°C, 10 min at 95 °C, 40 cycles of 15 s at 95 °C for denaturation and 60 s at 60 °C.

### Hemagglutination inhibition (HI)

Paired serum samples collected during acute and convalescent phase of infection from animals (n=20) from Farm 2, were used to determine seroconversion to HPAI H5N1 virus using the HI test. Serum HI activity was determined using BPL inactivated A/Tk/IN/3707/22 antigen (clade 2.3.4.4b), as described previously. HI titers are expressed as log2 values, with 1 log2 being the minimum titer considered positive.

### Virus isolation

Virus isolation was performed in pooled milk samples from Farms 1 and 2. Approximately 5 ml of milk from individual animals were pooled and a total of 50 ml of pooled milk were centrifuged at 1,700 x g for 10 min at 4°C. The supernatant was discarded, and the pellet was resuspended in 5 ml of sterile PBS-BSA (1%) followed by centrifugation at 1,700 x g for 10 min at 4°C. The wash step was repeated one more time and the final pellet was resuspended in 1 ml PBS-BSA (1%). Virus isolation was conducted in bovine uterine epithelial cells (CAL-1, developed in house at the Virology Laboratory at AHDC) cultured in minimal essential medium (MEM, Corning Inc., Corning, NY) supplemented with 10% fetal bovine serum (FBS) and penicillin-streptomycin (Thermo Fisher Scientific, Waltham, MA; 10 U.mL^−1^and 100 µg.mL^−1^, respectively). Cells were cultured in T25 flasks and inoculated with 1 mL of the milk pellet resuspension from infected cows and incubated at 37 °C for 1 hour (adsorption). The inoculum was then removed, and cells were washed once with phosphate buffered saline and replenished with 1 mL complete growth media (MEM 10% FBS). Cells were monitored daily for the development of cytopathic effects (CPE) including cell swelling, rounding and detachment. When the CPE reached 70-80%, infected cells were harvested, and cell suspensions were collected after three freeze-thaw cycles. The identity of the isolated virus was confirmed by rRT-PCR, an immunofluorescence assay (IFA) using anti-nucleoprotein mouse monoclonal antibody (HB65, ATCC, H16-L10-4R5) and whole genome sequencing. Cell nuclei were stained with 4′,6-diamidino-2-phenylindole (DAPI) (ThermoFisher Scientific, 62248).

### Virus titrations

The infectious viral loads in milk and tissues of infected animals was quantified by viral titrations. For this, serial 10-fold dilutions of rRT-PCR positive milk samples and tissue homogenates were prepared in MEM and inoculated into CAL-1 cells in 96-well plates. Each dilution was inoculated in quadruplicate wells. At 48h post-inoculation, culture supernatant was aspirated, and cells were fixed with 3.7% formaldehyde solution for 30 min at RT and subjected to IFA using the anti-NP (HB65) mouse monoclonal antibody. Virus titers were determined using end-point dilutions and the Spearman and Karber’s method and expressed as TCID_50_.mL^−1^.

### Microscopic changes, in situ hybridization (ISH) and immunohistochemistry (IHC)

A total of 25 tissue samples from four dairy cattle and 12 tissues from one domestic cat were collected and fixed in formalin. This formalin fixed paraffin embedded (FFPE) tissues were sectioned at 3 µm thickness, stained with hematoxylin and eosin (H&E), and examined for histological changes. To determine the virus tropism and tissue distribution in dairy cattle and cat affected with HPAI H5N1, we performed ISH and IHC on FFPE tissues as previously described^6^. Briefly, tissue sections were deparaffinized with xylene, washed with absolute ethanol, blocked with peroxidase followed by antigen retrieval for one hour. For the ISH the V-InfluenzaA-H5N8-M2M1 probe (Advanced Cell Diagnostics, Inc., Newark, CA) which targets H5Nx clade 2.3.4.4b viruses and the RNAScope HD 2.5 assay were used as per manufacturer’s instructions. ISH signals were amplified with multiple amplifiers conjugated with alkaline phosphatase enzymes and finally incubated with red substrate at room temperature for 10 minutes and counterstained with hematoxylin. Immunohistochemistry was performed at the University of Georgia Veterinary Diagnostic laboratory and the USDA-ARS Southeast Poultry Research Laboratory following standard diagnostic IHC procedure. Specifically, tissue sections were treated with Proteinase K for 5 min for antigen retrieval and monoclonal antibody (Meridian Bioscience, Catalog No. C65331M) to Influenza A virus M-gene was used at 1:100 dilution for 1 hour. All the slides were counterstained with hematoxylin, scanned at 40X resolution and the digital slides were examined for virus tropism and tissue distribution.

### Viral metagenomic sequencing

#### Sample Collection and Processing

Whole blood nasal swab samples were obtained from 10 cows from Farm 1 in Texas. Samples were submitted to the AHDC at Cornell University, on March 16, 2024. Upon receipt, metagenomic sequencing using the sequence-independent, single-primer amplification (SISPA) procedure, the Oxford Nanopore sequencing chemistry and GridION sequencing platform were performed as described below.

#### Nucleic Acid (NA) Extraction, Library Preparation and Sequencing

Nucleic acid (NA) extraction was performed in 190 µl from each sample using the QIAamp MinElute Virus Spin Kit (Qiagen). Prior to NA extraction samples were subjected to an enzymatic cocktail treatment composed of 10X DNase 1 buffer, DNAse 1, Turbo DNAse, RNase Cocktail (ThermoFisher Scientific), Baseline ZERO DNAse (Lucigen), Benzonase (Sigma-aldrich) and RNase ONE Ribonuclease (Promega) to deplete host and bacterial nucleic acid. Purified NA was subjected to SISPA, modified from a previously reported protocol ^47^ Briefly, 11 μL of nucleic acid was used in a reverse transcription reaction with 100 pmol of primer FR20RV-12N (5’-GCCGGAGCTCTGCAGATATCNNNNNNNNNNNN-3’) using SuperScript IV reverse transcriptase (Thermo Fisher Scientific), followed by second-strand synthesis using the Klenow Fragment of DNA polymerase (NEB) with primer FR20RV-12N at 10 pmol. After purification using Agencourt AMPure XP beads (Beckman Coulter), SISPA PCR amplification was conducted with TaKaRa Taq DNA Polymerase (Takara) using the primer FR20RV (5’-GCCGGAGCTCTGCAGATATC-3’) at 10 pmol. SISPA products were converted into sequencing libraries using the ligation sequencing kit (SQK-LSK109) and Native Barcoding Kit 96 V1 for multiplex sequencing. Sequencing was performed on the FLO-MIN106 MinION flow cell r9.4.1 using the GridION Sequencer (Oxford Nanopore Technologies). A 24-hour sequencing run was conducted, with fastq generation performed by the GridION using high accuracy base calling. Settings were adjusted to accommodate barcodes at both ends and filter mid-strand barcodes. Fastq reads were then filtered by size and quality using Nanofilt^48^ and classified using Kraken version 2.1.0^49^ followed by Bracken^50^.

### Targeted Influenza A Sequencing

Samples that tested positive for HPAI H5N1 and had Ct values <30 were subjected to targeted influenza A sequencing at the Animal Health Diagnostic Center at Cornell University (Cornell AHDC) and the Ohio Animal Disease Diagnostic Laboratory (Ohio ADDL). The set of 107 samples included samples from Farm 1, n=19; Farm 2, n=33; Farm 3, n=54; and Farm 7, n=1. A complete metadata table with details on this set of samples is provided in **Supplementary Data Table 1**. Initial targeted sequencing attempts on milk samples at Cornell AHDC utilizing high-throughput diagnostic extraction methods^6^, were unsuccessful in obtaining whole influenza A genome sequences despite the utilization of samples with low cycle threshold (Ct) values. To overcome this limitation up to 50 ml of each milk sample were pelleted at 1,770 x g for 15 min at 4°C. The pellets were washed two times in PBS as described above and resuspended in 1 ml of PBS-BSA. The resuspended pellet was then diluted 1:5 or 1:10 in PBS and 200 µl of this dilution were used for extraction with the Indical IndiMag Pathogen kit (INDICAL Bioscience) on the KingFisher Flex extractor (Thermo Fisher, Waltham, MA, USA). Whole influenza A virus genome sequences were generated using the MBTuni-12 and MBtuni-13 M-RT-PCR methods^51^. Sequencing libraries were generated using the Native Barcoding Kit, EXP-NBD196, Ligation Sequencing Kit, SQK-SQK109 (Oxford Nanopore Technologies [ONT]), and sequenced on a FLO-MIN106 MinION flow cell r9.4.1 using the GriION platform.

Additionally, 31 samples from Farm 3 were subjected to target influenza A sequencing at the OADDL using the Illumina DNA Prep Kit and the Nextera DNA CD Indexes. Paired-end sequencing was performed on an Illumina MiSeq platform using the MiSeq Reagent Kit V3 (Illumina) with 2×250 base pair chemistry.

### Sequence analysis and mutational profiling

Sequencing data generated by the GridION platform underwent high-accuracy basecalling and demultiplexing of barcodes. Settings were configured to require barcodes at both ends and to exclude reads with mid-read barcodes. The Nanofilt software version 2.8.0^48^ was employed to filter sequences based on quality thresholds. Reads with a quality score below 12 and those shorter than 600 base pairs were removed from further analysis. Filtered reads were aligned to a reference genome download from GenBank (A/Gallus/gallus_domesticus/Sonora/CPA-18486-23/2023/H5N1, NCBI accession numbers OR801090.1 through OR801097.1) using Minialign software version 0.4.4 (https://github.com/ocxtal/minialign). Consensus sequences were generated using Medaka software version 1.4.3 with medaka_haploid_variant and medaka_consensus programs for polishing (https://github.com/nanoporetech/medaka). Sequences with a read depth greater than 20 and a quality score exceeding 20 were retained. Analysis of Illumina MiSeq data was performed by trimming the reads with Trimmomatic version 0.39^52^, and aligning, calling variants and generating consensus sequences with Snippy version 4.6.0 ( https://github.com/tseemann/snippy). Genome sequences were annotated using Prokka software version 1.14.5 to identify genetic features and functional elements^53^. The GenoFLU tool version 1.03 assessed potential reassortment events within the viral genome (https://github.com/USDA-VS/GenoFLU). Genome alignments, mutations, SNPs, and annotation data were visualized using Geneious Prime software (version 1014.0.5). The FluServer tool, available through GISAID EpiFlu, was utilized to interpret the effects of mutations identified in the sequences, leveraging previously published data (https://flusurver.bii.a-star.edu.sg/). Other mutation data was visualized using protein consensus alignments in Geneious Prime software.

### Phylogenomic and Phylogeographic Analysis

The dataset consisted of HPAI H5N1 clade 2.3.4.4b genomes from samples collected between January 2023 and March 2024 in the American continent, downloaded from GISAID Epiflu database^24^, and 91 complete genomes from the present study, that includes 50 genomes obtained from raw sequencing data, combined with another 41 complete genomes curated from the GISAID database that were obtained from the farms in our study (Farm 1, n=11; Farm 4, n=3; Farm 5, n=1; Farm 6, n=4, Farm 7, n=5; Farm 8, n=6, and Farm 9, n=11). The genomes generated in this study are deposited in GISAID database (**Supplementary Data Table 5),** and raw reads are available in the Sequence Read Archive (SRA) under BioProject accession number PRJNA1114404. Phylogenetic analyses were performed by using procedures implemented in Nextstrain^54^. Briefly, multiple sequence alignment was performed using Nextalign; maximum likelihood tree was inferred using IQ-TREE through Augur tool kit and data visualization through Auspice. The potential transmission networks between farms were inferred using the PB2 gene sequences in PopART package v1.7.2 using median joining tree method with an epsilon of zero^55^.

---

**Extended Data Fig. 1.**
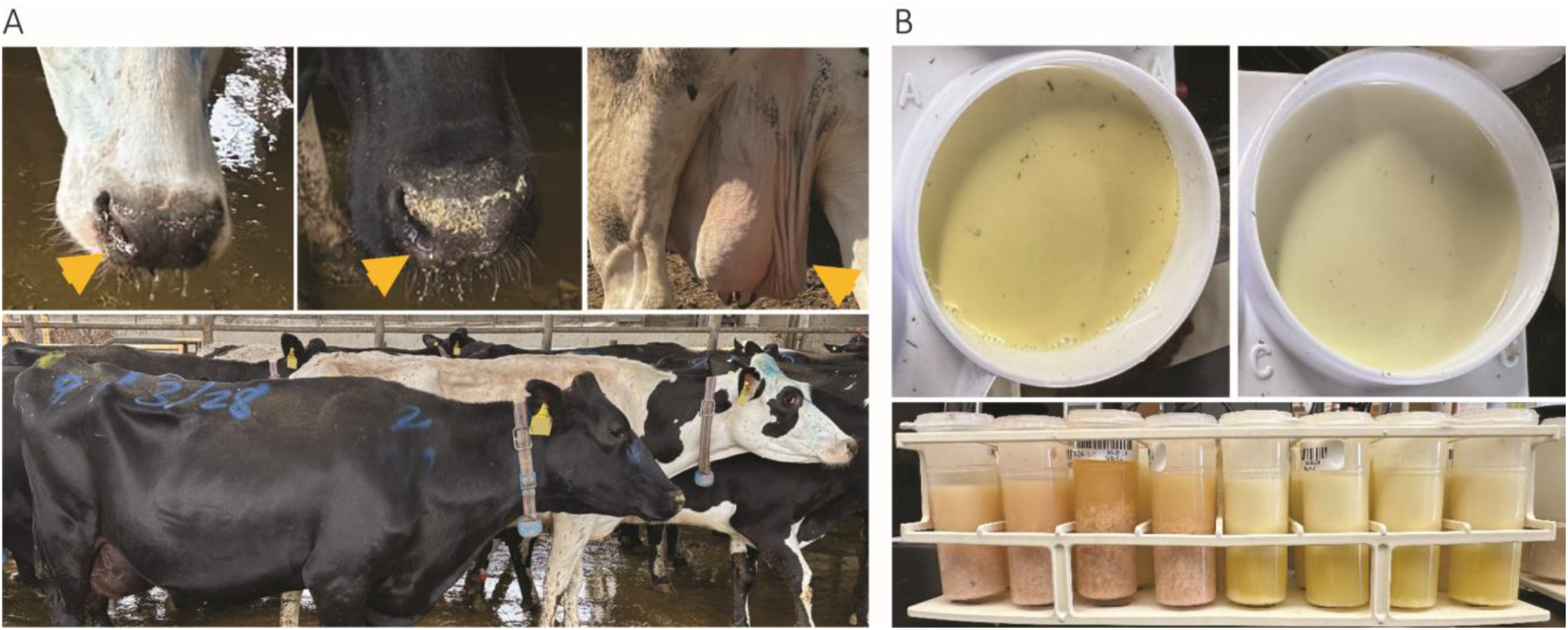
Clinical presentation of HPAI H5N1 infection in dairy cattle. (A) Infected animals presenting clear nasal discharge and involution of the mammary gland/udder (gold arrowheads, top images) and depression (bottom images). (B) Milk from HPAI H5N1 infected animals presenting yellowish colostrum-like color and appearance (top panels) or coloration varying from yellowish to pink/brown color. Curdling of milk visible in some samples.

**Extended Data Fig. 2.**
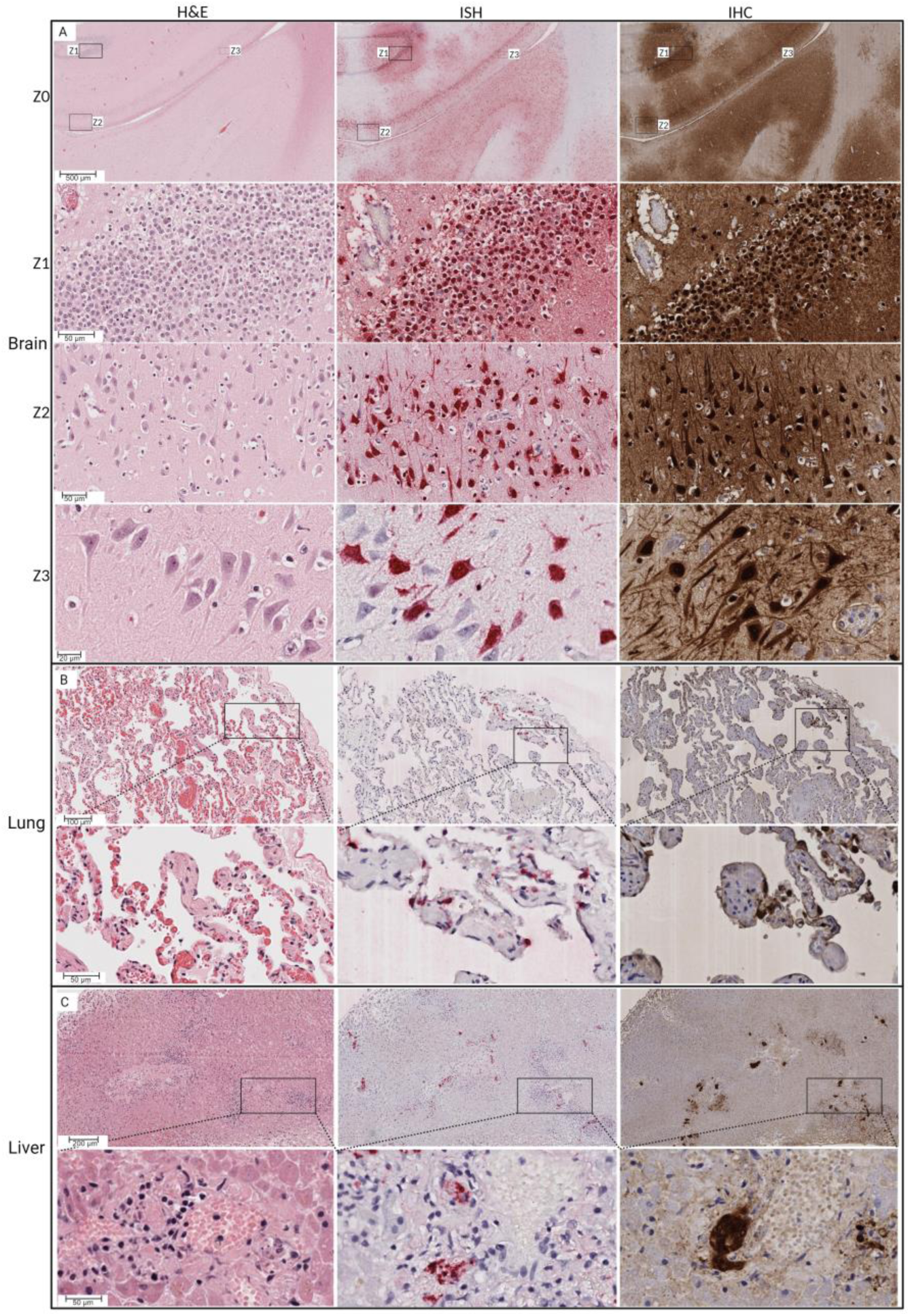
Highly pathogenic avian influenza virus H5N1 detection in cat tissue. Hematoxylin and eosin (H&E) staining (left panels) showing; A) multifocal area of perivascular cuffing, vascular congestion, and perivascular edema (Z0), neuronal swelling and neuronal necrosis and perivascular edema in brain (Z1, Z2 and Z3) B) pulmonary edema with strands of fibrin, thickened alveolar septa and intraepithelial lymphocytes, alveolar capillary congestion C) single cell necrosis and hemorrhage in liver. In situ hybridization (ISH) (middle panels) targeting Influenza A virus (Matrix gene) showing A) multifocal areas with extensive viral RNA (Z0), in neurons and glial cells within the granular layer and nuclear and intracytoplasmic viral RNA in neuronal soma, axon, and vascular endothelial cells in brain (Z1, Z2 and Z3), B) viral RNA in bronchiolar epithelial cells and type II pneumocytes, C) viral RNA in resident sinusoidal Kupffer cells and vascular endothelial cells. Immunohistochemistry (IHC) (right panels) targeting Influenza A virus M gene showing immunolabeling of A) multifocal areas of immunolabeling (Z0), intracytoplasmic immunolabeling of viral antigen in neuronal soma and axons within granular layer in brain (Z1, Z2 and Z3) B) bronchiolar epithelial cells and type II pneumocytes in lung, C) vascular endothelial cells and resident sinusoidal Kupffer cells.

**Extended Data Fig. 3.**
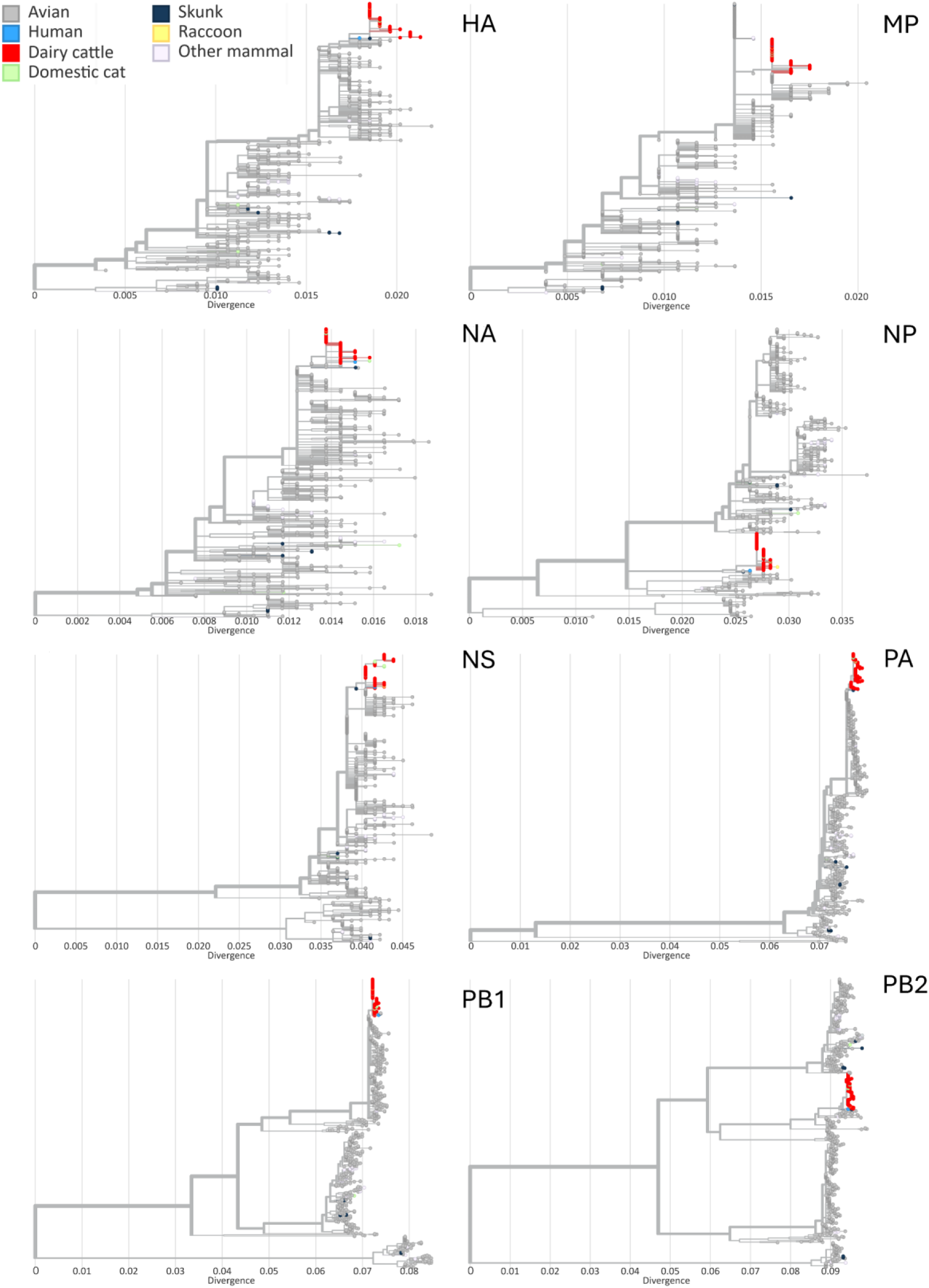
Phylogenetic trees of each genome segment, comprising sequences of 91 samples described in this study and sequences of 648 samples from the American continent, collected between January 2023 and March 2024, available in the GISAID EpiFlu database.

**Extended data Fig. 4.**
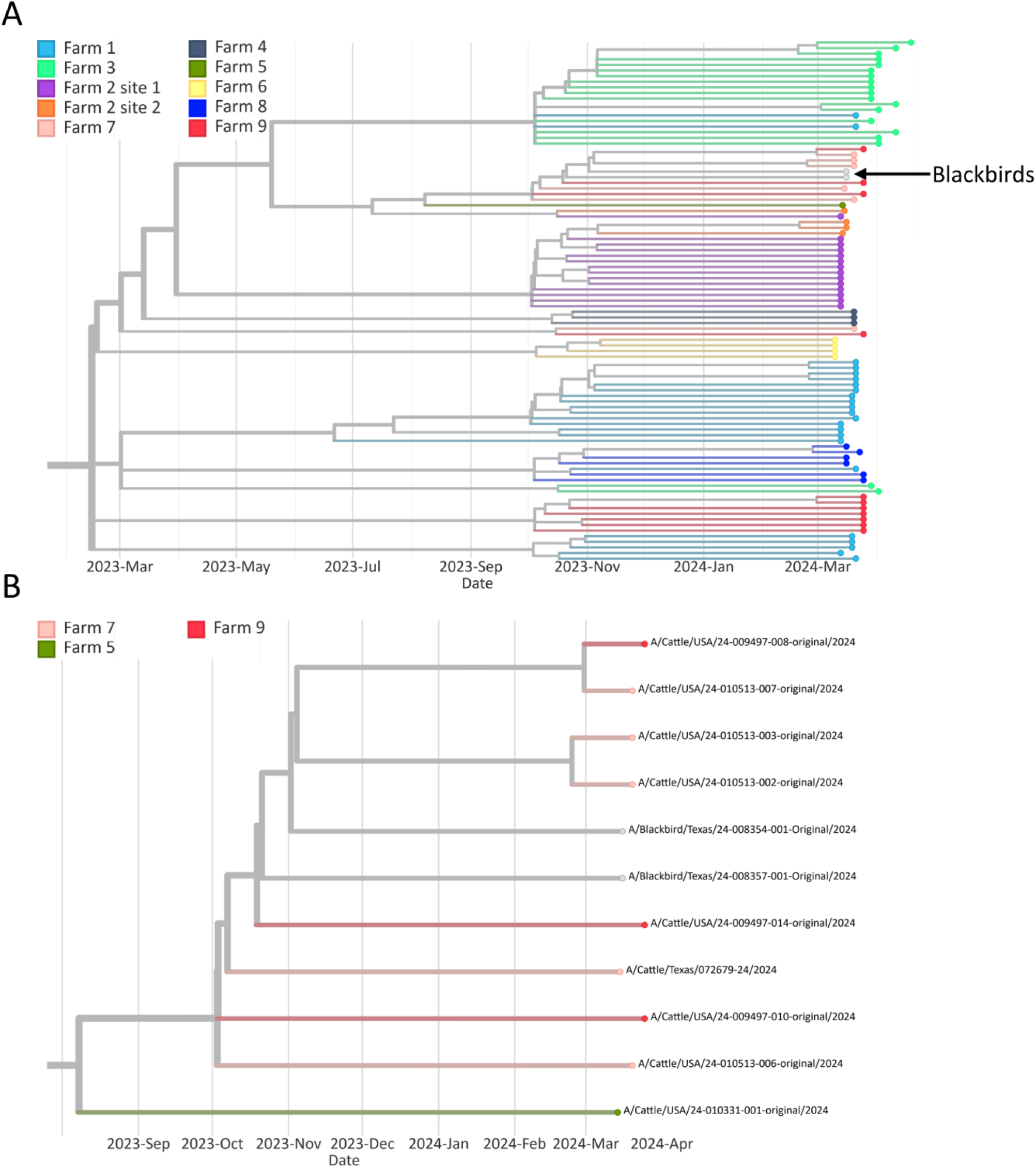
Wild bird sequences are related to sequences farm in affected dairy farms. (A) Genetic relationship of HPAI H5N1 sequences recovered from blackbirds with sequences recovered from cattle in Farm 7 and 9. Nodes are colored by premise and all the samples collected in the referred farm are highlighted. (B) Detailed/zoom in view of the sequence clusters containing samples from Farm 7, Farm 9 and the blackbirds collected at 8-12 Km from Farm 7.

**Extended data Fig. 5.**
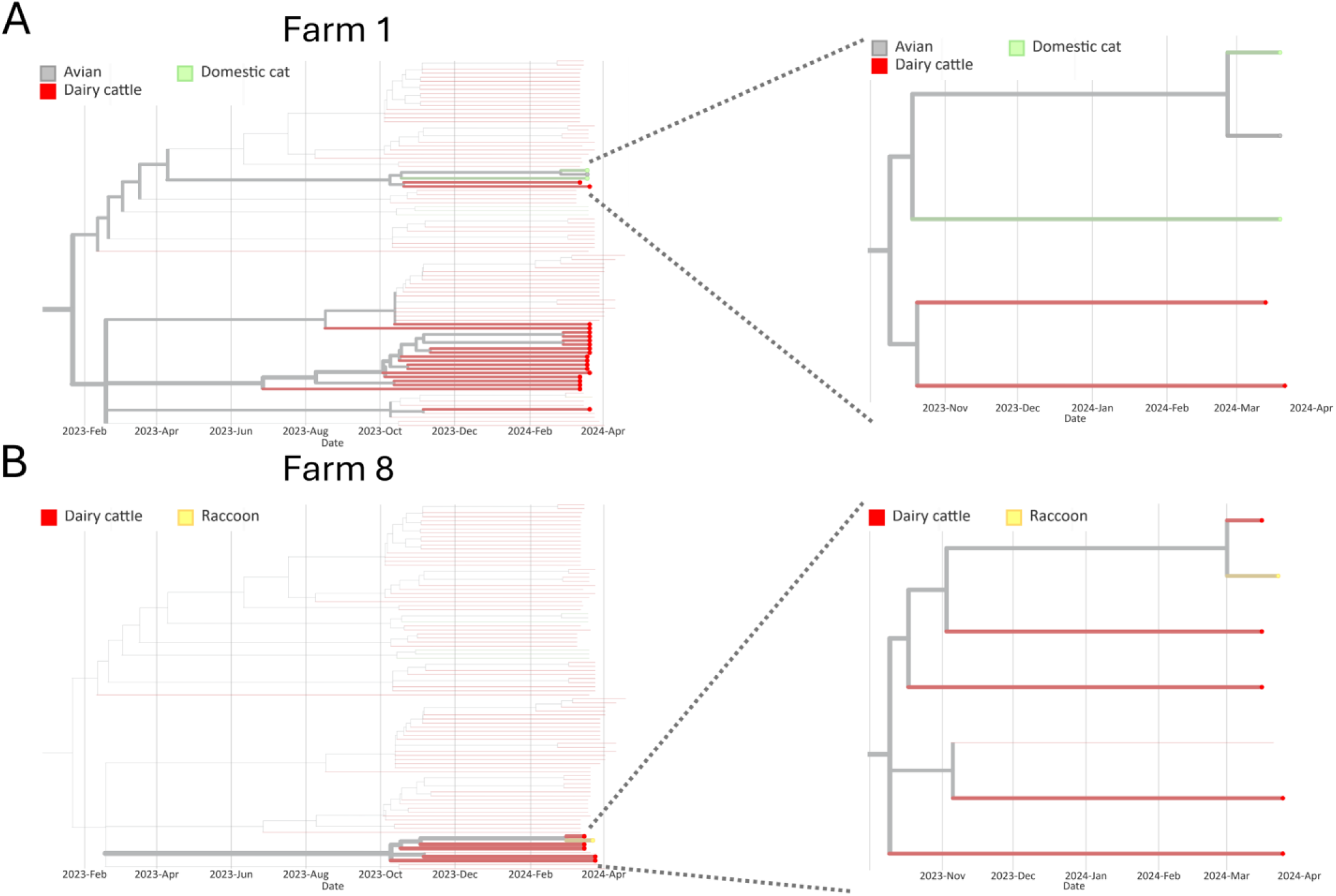
Evidence of interspecies transmission of HPAI H5N1 in (A) Farm 1 and (B) Farm 9. Nodes are colored by host and all the samples collected in the referred farm are highlighted. Panels on the right are a detailed view of the clusters containing more than one host species.

## Notes

### Competing Interest Statement

The authors have declared no competing interest.

